# PEBP1 is a candidate biomarker of response to the PP2A inhibitor LB-100

**DOI:** 10.1101/2025.02.25.640036

**Authors:** Matheus Henrique Dias, Chrysa Papagianni, Rene Bernards

## Abstract

We recently proposed an approach for cancer therapy involving a “paradoxical” activation of oncogenic signaling combined with the inhibition of stress responses. However, as with any other treatment, resistance can also emerge with hyperactivation therapy. In this study, we explored how cancer cells can acquire resistance to a drug that hyperactivates oncogenic signaling using the Protein Phosphatase 2A (PP2A) LB-100 as an example.

Our findings indicated that PEBP1 depletion confers resistance to LB-100 in different cancer models. Mechanistically, resistance is mediated by a reduced conversion of the prodrug LB-100 into the active metabolite endothall in the absence of PEBP1. Our data are compatible with a model in which PEBP1 is a hydrolase that can convert the prodrug LB-100 into the active endothall.

## Introduction

Colorectal (CRC) and pancreatic (PC) cancer are two prevalent and lethal adult malignancies accounting for nearly 900,000 and 400,000 deaths annually worldwide [1–3]. Both diseases have a low survival rate and impact individuals of all ages and genders [4–6]. Treatment for CRC and PC typically involves surgery and chemotherapy, but unfortunately, the long-term survival of the patients remains dismal. The prognosis is affected by a number of factors, such as the location of the malignancy, and the time of the disease discovery [2]. Both types of tumors usually remain asymptomatic during their formation and progression, and they are often diagnosed when cancer is already aggressive and metastatic [2, 7]. Surgery is beneficial for non-metastatic tumors, while chemotherapy is also used for metastatic cases, inhibiting the proliferation of rapidly dividing cells, and leading to apoptosis. Unfortunately, certain normal cell types also grow at high rates and are targeted during treatment, including hair follicles, blood, stem, and gastrointestinal tract cells, resulting in significant adverse effects [6].

The class of targeted therapies employs small molecule inhibitors, as well as monoclonal antibodies to suppress signaling transduction, aiming to kill cancer cells [6, 8, 9]. Moreover, these targeting compounds can be combined with chemotherapy, to increase the antitumor effect. Multiple targeted drugs are currently used in cancer therapy improving the overall prognosis in many cases [6].

Targeted therapy has revolutionized the therapeutic possibilities for some cancers since it increased overall long-term survival with reduced side effects. It has also led to a deeper understanding of cancer biology, allowing for more personalized and effective treatments based on the patient’s genetic background [10]. However, the major obstacle faced by cancer therapies is resistance [11], which can be classified into two main categories: primary or intrinsic and secondary or acquired. In primary resistance, the tumor is refractory to the treatment, and it is typically caused by the presence of feedback loops. The secondary resistance follows an initial response to the medication and is the outcome of the tumor cells’ evolution [10–12]. Resistance precludes the better efficacy of targeted drugs and resistant tumors often show aggressive phenotypes and lack alternatives for treatment [10]. Therefore, identifying ways to circumvent the impact of drug resistance on the disease’s prognosis is urgently needed.

Cancer cells can remarkably switch between mechanisms to achieve the most efficient phenotype, enabling them to proliferate, evade the immune system and resist cell death [13, 14]. Enhanced mitogenic signaling induced by somatic mutations is a hallmark of cancer, but this is not enough to lead to tumorigenesis [14]. For instance, the DNA damage associated with the increase in oncogenic messages disrupts cellular balance [15] and induces apoptosis [14]. During the processes of oncogenesis, the cells activate stress response pathways to counterbalance the excessive mitogenic signaling, maintain homeostasis and cancer cell viability [13, 16–18]. Therefore, in cancer progression, the oncogenic overactivation reaches optimal levels that are not necessarily the highest, since this comes at the expense of increased cellular stress [13, 19].

The concept that increased oncogenic signaling alone is insufficient to cause cancer and can only induce the disease when paired with the activation of stress response factors is well-established. For instance, overexpression of myc induces replication stress and combined with the pharmacological inhibition of anti-apoptotic proteins, such as Bcl-2, leads to apoptosis in cancer cells [14, 20, 21]. Additionally, Dias et al (2019) proposed the synthetic lethality between FGF2 and stress-targeted therapies. FGF2 administration further increases the oncogenic signaling, replication, and proteotoxic stress and sensitizes cancer cells to stress response inhibitors, including checkpoint or proteasomal inhibitors [16]. Another study by Cararo-Lopes et al (2021) supported that Ras mitogenic stimulation induces the overloading of stress response pathways, leading to autophagy of “HPV-infected” keratinocytes [22]. In light of the aforementioned findings, Dias and Bernards (2021) proposed “paradoxical overactivation” as a new therapeutic strategy for cancer. This approach involves the hyperactivation of the oncogenic pathways as a tool to disrupt cellular equilibrium and induce cellular stress. The combination of oncogenic activators with stress response inhibitors can be highly toxic for cancer cells [13].

Building upon this concept, we recently proposed the use of a small molecule activator of the oncogenic cascades to sensitize cancer cells to stress-targeted inhibitors [23]. The small molecule in question is LB-100 (3-(4-methyl piperazine-1-carbonyl)-7-oxabicyclo [2.2.1] heptane-2-carboxylic acid) which inhibits the protein phosphatase 2A (PP2A) [24, 25]. LB-100 demonstrated chemo-sensitizing effects in several cancer models, such as glioblastoma, sarcoma, breast, pancreatic, and ovarian cancer, both *in vitro* and *in vivo* [24, 26]. A number of mechanisms by which LB-100 exerts its cell toxicity have been proposed. For instance, recent data has shown that it can lead the cells to apoptosis through decreased DNA damage repair [24], G2/M cell-cycle arrest [24, 26], increased drug penetration, and induction of mitotic catastrophe [24]. Importantly, LB-100 has already undergone a phase I clinical trial, showing antitumor activity and moderated associated toxicity, rendering it a promising candidate for cancer therapy, either alone or in combination with other treatments [27].

PP2A is a serine/threonine phosphatase and plays a pivotal role in signaling transduction [24]. PP2A dephosphorylates factors of the Wnt/ β-catenin, PI3K/Akt, and MAPK pathways, including Akt, GSK3-β, Erk, and MEK [28, 29]. It, additionally, dephosphorylates anti-apoptotic proteins, such as Bcl-2 and Jak/Stat cascade components [24, 28], and regulates mitotic proteins (Figure 1) [23]. Based on its function, it has a significant role in cellular proliferation, division, and death, and in general in cellular homeostasis [29].

**Figure 1.**
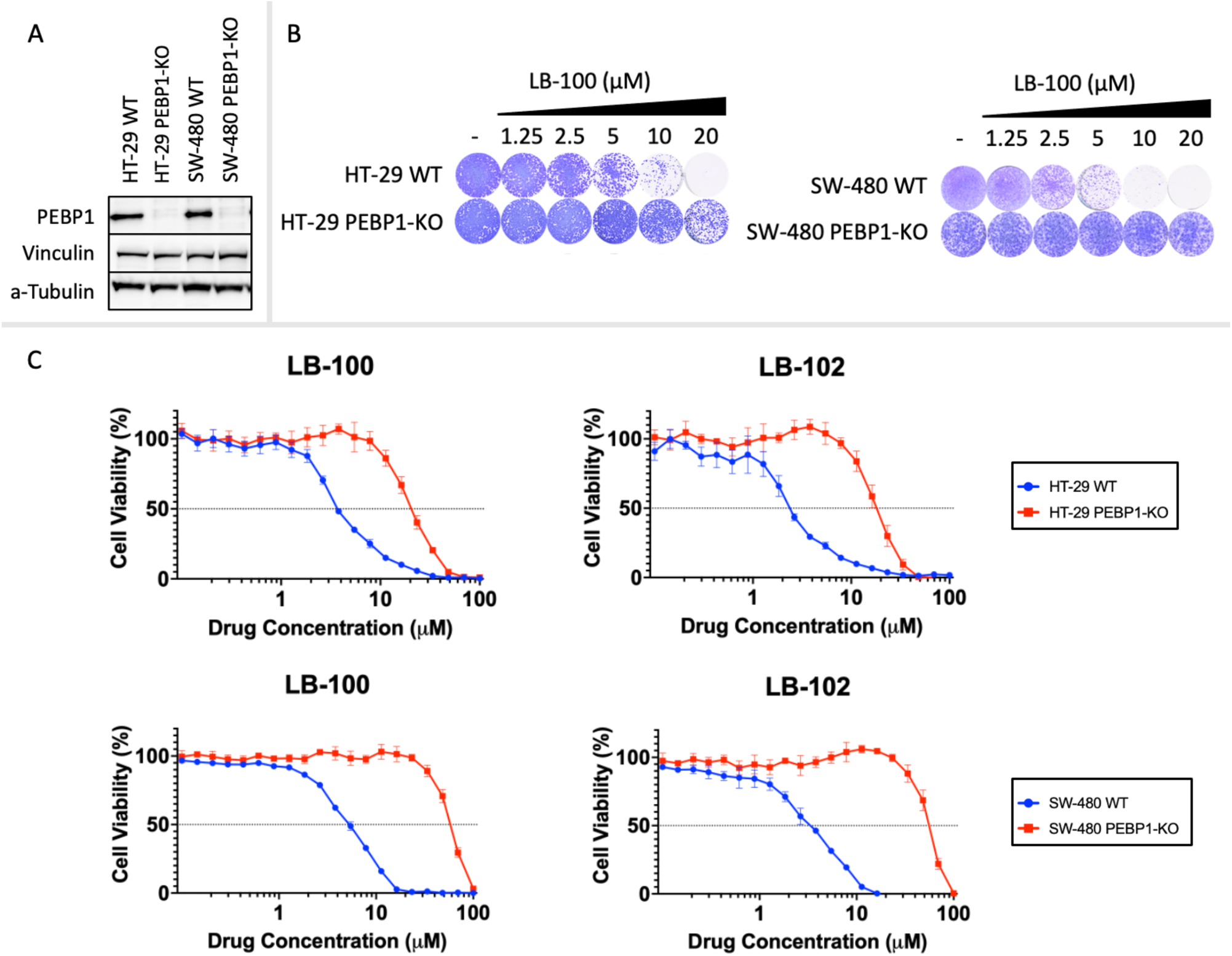
(A) Representative Western blot analysis of PEBP1 to validate the CRISPR-based PEBP1-knockout (KO) in the HT-29 and SW-480 cell lines. The wild-type [43] cells were used as a control for the expression of PEBP1. Vinculin and a-Tubulin were used as housekeeping proteins. (B) Long-term viability assays of HT-29 and SW-480 WT and PEBP1-KO cells under treatment with LB-100. The assay was terminated when the untreated group reached confluence. (C) Dose-response curves of HT-29 and SW-480 WT and PEBP1-KO cells under treatment with the water-soluble compound LB-100 and the lipid-soluble compound LB-102. The values were normalized to PAO and DMSO, which were used as a positive control (0% cell viability) and negative control (100% cell viability), respectively. The mean of the values and their SD are illustrated.

PP2A controls a myriad of pathways that are often dysregulated in cancer and therefore it is a tumor suppressor protein [24, 29]. Because of its inhibitory effect on oncogenic pathways, inhibition of its phosphatase activity should lead to oncogenic signaling overactivation and a consequent increase in stress responses [29, 30]. In that concept, LB-100 suppresses PP2A, subsequently overactivates oncogenic signaling, and renders cancer cells reliant on stress responses. Hence, a promising therapeutic combination could pair LB-100 with a stress response inhibitor. A screening of compounds, targeting stress response pathways frequently associated with oncogenic phenotypes, revealed that LB-100 increased the toxicity of the WEE1 inhibitor, adavosertib, against cancer cells [23].

Despite the mechanistic rationale and the efficacy observed for this combination, resistance is likely a major hindrance to any prospective cancer treatment. Therefore, exploring potential mechanisms of resistance to this combination, a genome-wide CRISPR-knockout screen identified genes whose down-modulation could result in resistance to LB-100. gRNAs targeting proto-oncogenes from the Wnt and MAPK pathways, including β-catenin, BCL9L, MAPK14, and MAPK1/ERK2, were enriched in the cells treated with LB-100. This is in agreement with the notion that if LB-100 overactivates oncogenic signaling, attenuation of this signaling should reduce the toxicity of the drug [23]. Interestingly, PEBP1 (phosphatidylethanolamine-binding protein 1), a negative regulator of the MAPK pathway [31], was identified as the top hit on the screen, suggesting a key role of this protein in mediating LB-100 toxicity [23].

PEBP1 is a cytoplasmic protein that is physically linked with the cellular membrane by interacting directly with phosphatidylethanolamine (PE). The protein is expressed intracellularly and has been found in the brain, colon, lung, and pancreas, among other organs. It is a small (21-23 kDa) protein with a defined structure and has been associated with many critical cellular processes, by regulating various signaling pathways such as MAPK, NF-κB, GPCR, GSK3β, JAK/STAT, and PI3K/Akt/mTOR [31–33]. PEBP1 comprises two primary domains, the ligand binding pocket, and the protein-protein interaction domain, which enable it to interact with various ligands and proteins and perform its diverse functions [31].

The function of PEBP1, also known as RKIP (Raf kinase inhibitory protein), is mainly controlled by the protein kinase C (PKC) (Figure 2). PKC phosphorylates PEBP1 at serine 153 (S153), which serves as a switch regulating the interaction of PEBP1 with its targets. S153 phosphorylation enables PEBP1 to bind with the G protein-coupled receptor kinase 2 (GRK-2) and decreases PEBP1’s affinity to Raf kinase. The dimerization of PEBP1 with GRK-2 modulates the signaling of the GPCRs, disrupting receptor internalization, and resulting in the permanent activation of GPCRs and Raf. Conversely, when PEBP1 is unphosphorylated at S153, it interacts with Raf, governs its activation, and blocks the MAPK pathway, by preventing the activation of MEK1/2 and subsequently Erk1/2 and the MAP kinases. Moreover, PEBP1 can directly interact with various crucial components of the NF-κB pathway, forming a multicomponent inhibitory complex, and thereby influencing its activation. Additionally, it directly attaches to GSK-3, stabilizing it and blocking its p38 MAPK-mediated inhibitory phosphorylation. Furthermore, PEBP1 interacts with STAT3, leading to the disruption of STAT3 phosphorylation by c-Src. Lastly, it inhibits the PI3K/Akt/mTOR pathway by binding to and inhibiting Raf-1. Overall, PEBP1 is a multifunctional protein that plays a crucial role in regulating numerous signaling pathways and therefore cellular processes [31, 34–36].

**Figure 2.**
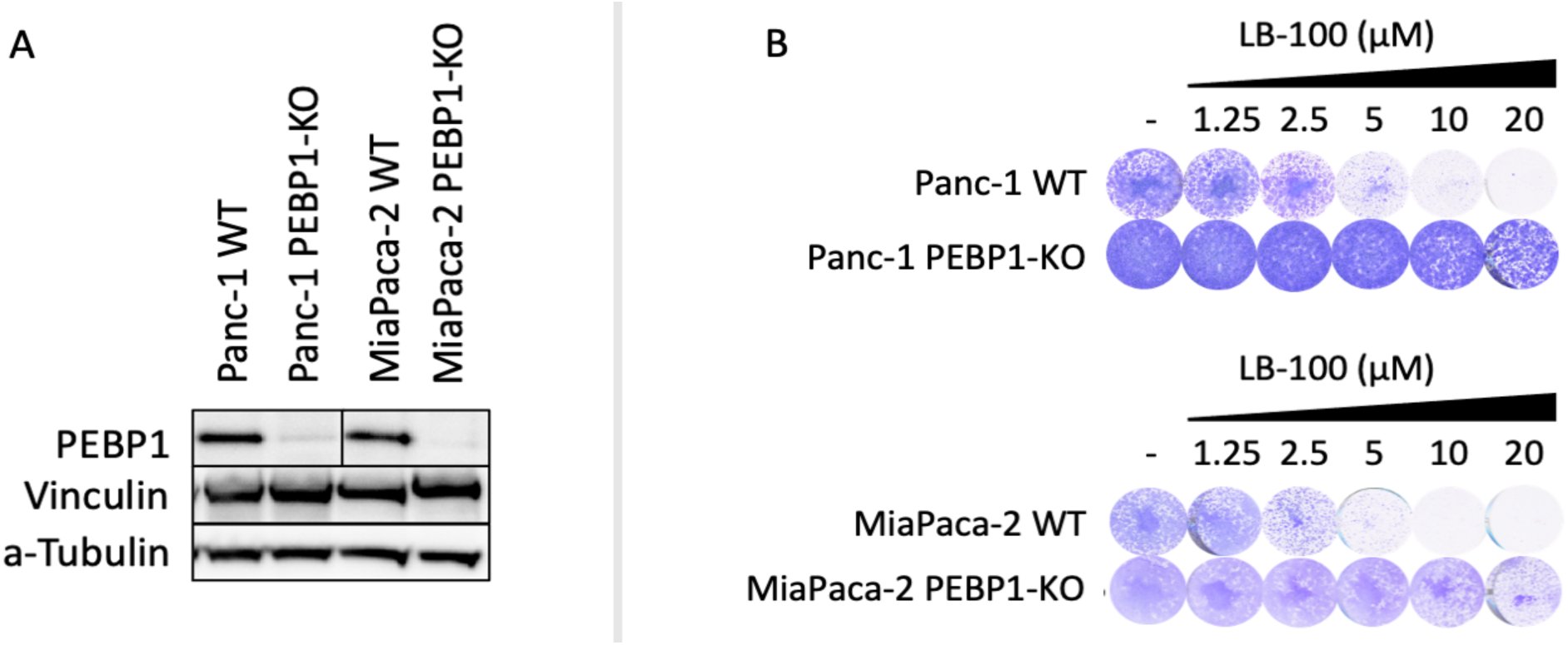
(A) Western blot analysis of PEBP1 to validate the CRISPR-based PEBP1-KO in the Panc-1, and MiaPaca-2 cell lines. The WT cells were used as a control for the expression of PEBP1. Vinculin and a-Tubulin were used as housekeeping proteins. (B) Long-term viability assays of Panc-1 and MiaPaca-2 WT and PEBP1-KO cells under treatment LB-100. The assay was terminated when the untreated group reached confluence.

The involvement of PEBP1 in numerous signaling pathways proposes that it has a significant role in cancer development, progression, and migration. It is commonly downregulated in the majority of cancer types, including but not limited to the colon, brain, kidney, and lung, and often absent in metastatic tumors [33, 35, 36]. Moreover, evidence has shown that low expression of PEBP1 is frequently associated with poor survival in renal, pancreatic, and colon cancer to name a few. The prognostic value of low PEBP1 expression is, in general, supported by the notion that PEBP1 is a tumor suppressor protein, as it acts in favor of mitogenic inhibition [33, 37]. Therefore, from that perspective the fact that its knock-out led to resistance to LB-100 [23] is counter-intuitive and deserves investigation.

In a previous study, Kazui et al (2016) suggested that PEBP1 can also function as a hydrolase and participate in prodrug metabolism [38]. Prodrugs are bioreversible compounds that undergo enzymatic or chemical processes to release an active metabolite with therapeutic potential [39, 40]. According to their findings, PEBP1 contributes to the conversion of the prodrug Prasugrel (a thienopyridine-type antiplatelet prodrug) into its active metabolite, thiolactone, and it is proposed as a dynamic protein, modulating not only intracellular signaling pathways but also drug metabolism [38].

Resistance is the Achilles heel of cancer therapies. Our recent data suggest that the depletion of PEBP1 or different oncogenic proteins may attenuate the toxicity of LB-100. In light of these findings, we endeavored to elucidate the mechanism behind LB-100 resistance. Specifically, we explored the resistance to LB-100 associated with low levels of PEBP1 and aimed to understand the underlying mechanism.

## 2. Materials and Methods

### Cell Lines and Cell Culture

The experiments were performed in HT-29 and SW-480 colorectal cancer, as well as in MiaPaca-2 and Panc-1 pancreatic cancer cell lines (Table 1). All cell lines were obtained by ATCC. The cells were cultured in Roswell Park Memorial Institute 1640 Medium (RPMI 1640, Gibco Fisher Scientific) supplemented with 10% Fetal Bovine Serum (FBS; Gibco Fisher Scientific), 0.5 U/mL penicillin and 50 μg/mL streptomycin (Sigma-Aldrich) at 37°C in a humidified, 5% CO_2_ atmosphere.

### Generation of PEBP1-KO clones

Generation of the PEBP1-knockout (KO) HT-29 and SW-480 cells was performed by transduction with the lentiCas9-Blast vector (Addgene, #52962) and a set of 4 synthetic oligos targeting PEBP1, obtained by Synthego. The transduction was conducted according to the manufacturer’s instructions. The edited HT-29 and SW-480 cells were selected with 10 µg/ml blasticidin (Invitrogen, #ant-bl-1) and the knock-out of PEBP1 was validated by Western blot. Blasticidin was also used as a positive control on non-transduced cells. The Panc-1 and MiaPaca-2 were transduced using a lentiCas9-Blast vector (Addgene, #52962), and a PEBP1-sgRNA in a lentiGuide-Puro vector (Addgene, #52963) with polybrene transfection reagent. The sequence of the sgRNA was GCATGTCACCTACGCCGGGG. Antibiotic selection with 10 µg/ml blasticidin and 2 µg/ml puromycin was performed, and the knock-out of PEBP1 was validated by Western blot. Blasticidin and puromycin were used as positive controls on non-transduced cells. Moreover, cells were transduced separately with the lentiCas9-Blast vector and the lentiGuide-Puro vector, and selection with blasticidin and puromycin was used as negative controls.

### Western Blot

The cells were washed with PBS and lysed with RIPA buffer (25mM Tris-HCl, pH 7.6, 150 mM NaCl, 1% NP-40, 1% sodium deoxycholate, 0.1% SDS) supplemented with a Complete Protease Inhibitor cocktail [41] and phosphatase inhibitor cocktails II and III (Sigma). Cellular fragments were removed by centrifugation at 14,000 x g at 4°C for 25 minutes. The quantification of protein concentration was performed using the PierceTM BCA Protein Assay Kit (Thermo Scientific) and BSA was used as a standard for the protocol. The samples were then normalized to a specific protein concentration, typically 0.7 µg/µl. Next, the cell lysates were mixed with NuPAGE™ LDS Sample Buffer 4X and Sample Reducing Agent 10X (Thermo Scientific) and were denatured by incubation samples at 100°C for 10 minutes. The proteins were separated in 4-12% polyacrylamide gels by SDS-PAGE (Bolt™ Bis-Tris Plus; Thermo Fisher Scientific) at 180 Volts for 45 minutes, transferred to nitrocellulose blotting membranes 0.45 µm (Amersham^TM^ Protran^TM^) at 350mA for 2 hours and blocked with 1% BSA and 1% non-fat dry milk diluted in TBS (10 mM Tris (pH 7.5) and 150 mM NaCl) with 0.1% Tween-20 (TBS-T) for 1 h at room temperature on a shaker. Subsequently, the membranes were probed with the primary antibodies diluted in 5% BSA at 4°C overnight. Next, they were washed 3 times with TBS-T buffer for 15 minutes and incubated with the secondary antibodies (HRP conjugated), diluted in 5% BSA, for 1 hour at room temperature. After the secondary antibody incubation, the membranes were washed 3 times TBS-T for 10 minutes. The membranes were stained with SuperSignal™ West Atto Ultimate Sensitivity Substrate (Thermo Scientific) and Clarity™ Western ECL Substrate (Bio-Rad) mixed in a ratio of 3:1. The protein bands were visualized by chemiluminescence using the ChemiDoc Imaging System (Bio-Rad). Vinculin, GAPDH, and a-tubulin were used as loading controls.

### Dose-Response Curves

Cells were placed with 20% confluence in black-walled 384 well plates (Greiner, #781091) and were incubated overnight. The next day, drugs were added using a Tecan D300e digital dispenser. Each drug or drug combination was tested in triplicates. 10 µM phenylarsine oxide (PAO, Sigma, #637-03-6) and 10 µM dimethyl sulfoxide (DMSO, Sigma, #67-68-5) were used as a positive control (0% cell viability) and negative control (100% cell viability), respectively. The plates were incubated for 5 days at 37°C in a humidified atmosphere. Resazurin (Sigma, #R7017) was added by a Thermo Multidrop Combi Dispenser and used to calculate cell viability. The cells were incubated for 1-4 hours depending on the cell type and the fluorescence was measured with EnVision (Perkin Elmer). The readings are normalized to the positive and negative control and the values are used to make dose-response curves.

### Colony Formation Assay

Cells were placed in 12-well plates (Greiner, #665180) at low density and were incubated overnight at 37°C in a humidified, 5% CO_2_ atmosphere. After cell attachment, the drugs were added using a Tecan D300e digital dispenser. The culture media and the drugs were refreshed every 2-3 days and the cells were fixed and stained when the controls were confluent. For the fixation/staining, a solution of 1% formaldehyde (Millipore, #104002), 1% methanol (Honeywell, #32213), and 0.05% crystal violet (Sigma, #HT90132) diluted in phosphate-buffered saline (PBS) was used. The plates were incubated for 20-30 minutes at RT. The plates were washed with ddH20, left to air-dry, and scanned using an EPSON V700/V750 scanner.

### Cell-free measurement of LB-100 conversion

BL21(DE3) cells were transduced to the LIC1.1 construct carrying the gene of the PEBP1 protein. The cells were lysed with 50 ml of lysis buffer (25 mM Tris pH 8.0, 200 mM NaCl, 1 mM TCEP) by sonication (3 minutes, 60% output, 10 sec on; 10 sec off; on ice). The cell debris and insoluble proteins were removed by centrifugation at 21,000 rpm at 4°C for 30 minutes. A gravity flow column with ∼3 ml nickel beads equilibrated with lysis buffer and the lysate was loaded onto it. After the lysate had been passed, the flow-through was collected and the beads were washed with 50 ml of wash buffer (lysis buffer + 20 mM imidazole). The PEBP1 protein was eluted in 5 fractions of elution buffer (lysis buffer + 200 mM imidazole) and the purification was validated by Western Blot. Next, the protein was injected into a S200 gel filtration column (120 ml volume), which had been equilibrated with buffer A (25 mM Tris pH 8.0, 200 mM NaCl, 1 mM TCEP), and fractions were collected using a BioRad NGC chromatography system. The purification was tested again with Western Blot. Finally, the peak fractions with PEBP1 were pooled and concentrated using an Amicon Ultra-15 concentrator.

To compare the amount of LB-100 and endothall in the presence or absence of recombinant PEBP1, the fraction from the endogenous enzyme purification was mixed with LB-100. The recombinant PEBP1 was used at 1 µg/ml and LB-100 was added at 50 µM in 0.1 M HEPES buffer in a total reaction volume of 200 µl. The reactions were performed in triplicates. The samples were LB-100 (5 minutes incubation), LB-100 + recombinant PEBP1 (5 minutes incubation), LB-100 (45 minutes incubation), and LB-100 + recombinant PEBP1 (45 minutes incubation). To stop the reactions, 600 µl of acetonitrile was added to each tube, making a total volume of 800 µl, with LB-100 diluted to 12.5 µM. After the precipitation, the samples were centrifuged at 14000 rpm at RT for 15 minutes and the supernatants were transferred in fresh tubes. Mass-spectrometry was performed to determine the concentration of the prodrug and the metabolite.

### 2.11 Compounds

LB-100 (#206834) was purchased by MedKoo. LB-102 and endothall were a kind gift of Lixte.

## Results

### PEBP1 depletion attenuates LB-100 toxicity

Our recent findings of the CRISPR-knockout screen indicate that suppression of PEBP1 attenuates the toxicity of LB-100 [23]. To validate the involvement of PEBP1 in the modulation of drug toxicity and later expand the insights into the underlying mechanisms, we generated CRISPR-based PEBP1-knockout cells. We used the CRC cell lines (HT-29 and SW-480) and the successful generation of the knockout clones was examined by Western blotting. The housekeeping proteins (Vinculin and a-Tubulin) demonstrated the accuracy of the protein loading. The analysis showed undetectable levels of the target protein in the knockout cell lines, while a distinct band was visible at the expected molecular weight in the control wild-type cells (Figure 1A). This observation confirms the efficacy of the gene editing strategy.

Subsequently to obtaining the knockout cells, we conducted long-term viability assays to verify the findings of the CRISPR-knockout screen. Our experiment was designed to compare the response of the wild-type and PEBP1-knockout cells to different concentrations of LB-100 (ranging from 1.25 – 20 µM). The results of the assay demonstrated that the knockout cell lines exhibited resistance to the treatment compared to the wild-type cells. The wild-type cells displayed a decline in cell viability under treatment, whereas the knockout cells maintained their growth at a similar rate to the untreated (Figure 1B). These data confirm the outcome of the screen and establish a correlation between reduced response to LB-100 and low expression levels of PEBP1.

Given the localization of PEBP1 in close proximity to the inner part of the cellular membrane and the fact that LB-100 is highly hydrophilic, we reasoned that PEBP1 could be involved in the intake of the drug. To explore this possibility, we performed viability assays using LB-100 and another PP2A inhibitor, known as LB-102 [25]. The primary difference between the two compounds lies in the presence of an ester group on LB-102, which renders it lipophilic. According to Lai et al (2018), both molecules have exhibited equivalent inhibitory effects on the PP2A activity and no toxicity to normal cells. Due to its lipid-soluble nature, LB-102 can be passively diffused across the cellular membrane [42] and therefore it was used as a control marker to test whether PEBP1 is involved in the internalization of the drug or its conversion. If so, the toxicity of LB-102 would not be affected by the depletion of PEBP1. Our data indicated that the deletion of PEBP1 resulted in similar levels of resistance to both LB-100 and LB-102, supporting that it is not implicated in the intake of the prodrug (Figure 1C). In summary, our findings suggest that PEBP1-depletion reduces LB-100 toxicity, and it is most probably not involved in its internalization.

### PEBP1 depletion is associated with resistance to LB-100 in different cancer cell models

Our findings revealed that decreased expression of PEBP1 is correlated with attenuated toxicity of LB-100. Based on that, we hypothesized that this phenomenon may not be restricted to specific cancer cell types but may be a general observation applicable to cell lines derived from other types of cancer. Therefore, we selected Panc-1 and MiaPaca-2 PC cell lines for further experimentation. The chosen cell lines derive from types of cancer in which PEBP1 downregulation is linked to poor patient prognosis and survival [33, 37] and they all harbor different mutations [23]. To perform our experiments, we generated Panc-1 and MiaPaca-2 PEBP1-depleted cell clones using CRISPR-based knock-out. Western blotting was employed to test the efficiency of the editing. Vinculin and a-Tubulin were used as loading controls. As observed for the CRC models, the results showed almost undetectable levels of PEBP1in the knock-out cells in comparison with the wild-types (Figure 2A), validating the efficiency of the genomic editing.

After obtaining the PEBP1-depleted cells, we performed long-term viability assays with increased concentrations of LB-100 (1.25 – 20 µM) on the Panc-1 and MiaPaca-2 cells. Our objective was to evaluate the influence of PEBP1 expression on the response to the treatment. The results exhibited that depletion of the protein reduced the sensitivity of the cells to LB-100 (Figure 2B). Collectively, our findings demonstrate a correlation between PEBP1-low levels and resistance to LB-100 in 4 cell lines from two distinct cancer types.

### Reduced inhibition of the total protein phosphorylation levels by PP2A in the absence of PEBP1

Our results indicated that PEBP1 modulates the effectivity of LB-100, but this membrane-bound protein was not involved in the internalization of the compound. As previously stated, LB-100 is an inhibitor of PP2A, a multifunctional phosphatase that regulates various signaling pathways by dephosphorylation [23, 24, 28–30]. Accordingly, inhibition of PP2A by LB-100 should increase the overall phosphorylation in cells. Based on the above, we asked whether low levels of PEBP1 would not only impact the toxicity of LB-100 but also its role in PP2A inhibition. If so, the absence of PEBP1 should reduce the overall phosphorylation induced by LB-100.

To explore the effect of PEBP1 on the mechanistic function of LB-100, we conducted a Western blot analysis to evaluate the phosphorylation levels of proteins in wild-type and PEBP1-knockout cells. We tested 3 different time points, 4, 8, and 24 hours of treatment. There was an untreated condition as a control group. We blotted for total phosphorylation of Threonine/Tyrosine and Serine/Threonine to cover a broad range of possible dephosphorylation activity. The images showed that for the wild-type cells, there was a significant increase in phosphorylation under treatment with LB-100 in comparison with the control. The highest effect was observed after 8 hours of treatment for all the cell lines tested. In contrast, mild or no increase in the phosphorylation levels were found in PEBP1-knockout cells after LB-100 treatment (Figure 3). These data indicate the essential role of PEBP1 in the inhibition of PP2A by LB-100, suggesting that PEBP1 is involved in the activation process of the prodrug.

**Figure 3.**
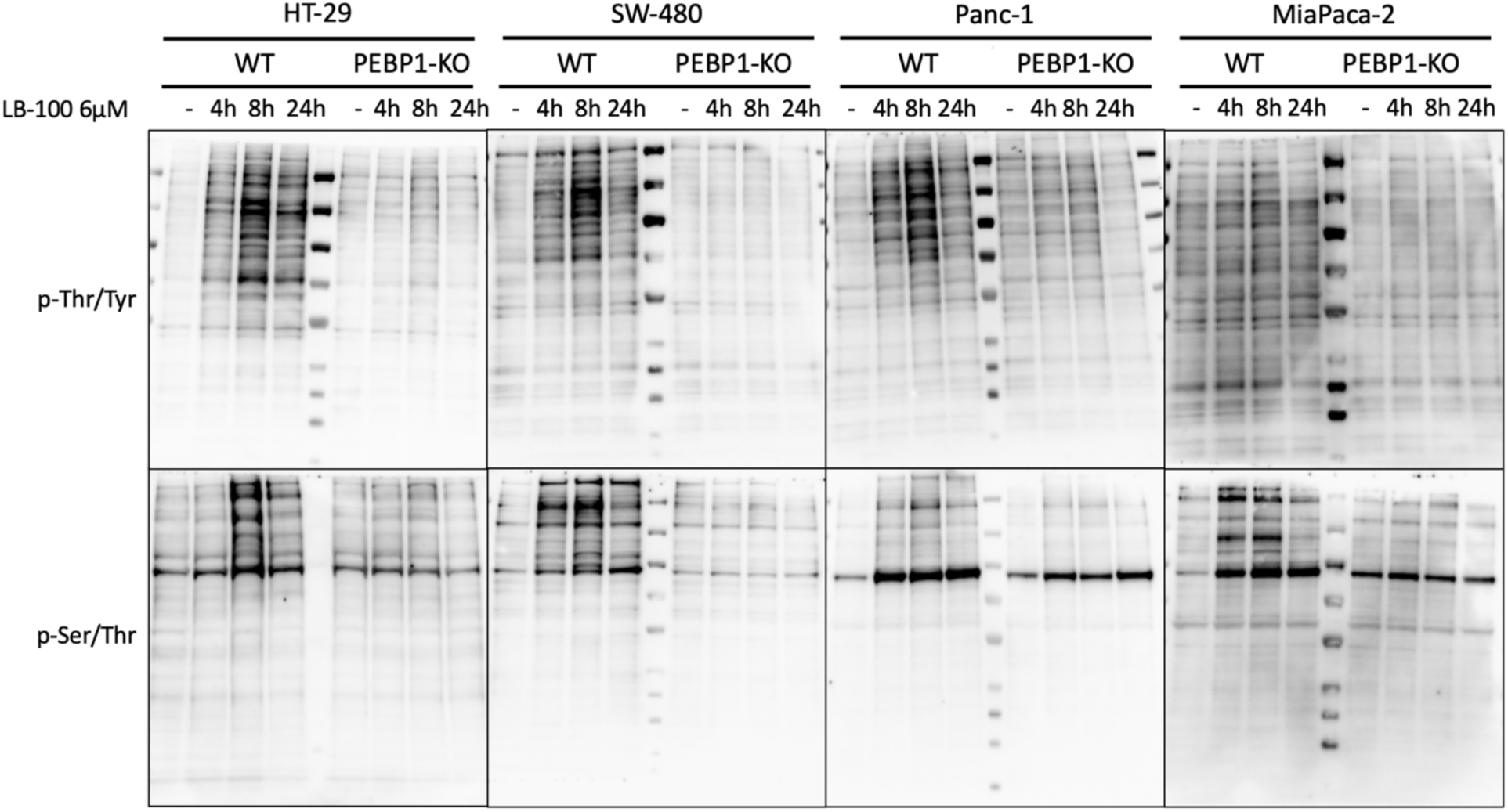
Western blot of the total phosphorylation levels of proteins in CRC and PC WT and PEBP1-KO cells. The total phosphorylation of Threonine/Tyrosine and Serine/Threonine were blotted after 3 different time points of treatment with 6 µM LB-100 (4, 8, and 24 hours). There was an untreated condition as a control group.

### PEBP1 is involved in the conversion of LB-100 to endothall

According to our observations, downregulated expression of PEBP1 led to resistance to LB-100 and influenced its inhibitory effects on PP2A in CRC and PC models. However, our experiments with LB-102 suggested that reduced drug internalization was not the cause of the resistance. Instead, they indicated that PEBP1 may be involved in modifying the drug after it enters the cell. Previous research by Kazui et al (2016) revealed that PEBP1 has an enzymatic function on Prasugrel. According to their findings, the protein of our interest diverts the prodrug Prasugrel into its active metabolite, thiolactone [38]. LB-100 is also classified as a prodrug since its active metabolite, endothall, is the one that exerts PP2A inhibition by interacting with its catalytic subunit [25], [44]. By combining the aforementioned insights, we asked whether the metabolic function of PEBP1 extends beyond Prasugrel and may be responsible for the LB-100 resistance in low levels of PEBP1.

To examine our hypothesis, we performed dose-response curves using CRC and PC wild-type and PEBP1-knockout cell lines treated with LB-100 and endothall. If the hypothesis was valid, the presence or absence of PEBP1 would not influence the effectiveness of endothall, but LB-100 would be more toxic when PEBP1 was present. In case PEBP1-depleted cells were also resistant to endothall, the protein would not be involved in the intermediary reaction of prodrug activation to the metabolite. The dose-response curves demonstrated that the knockout cell lines displayed resistance to the prodrug and not to the active metabolite (Figure 4A).

**Figure 4.**
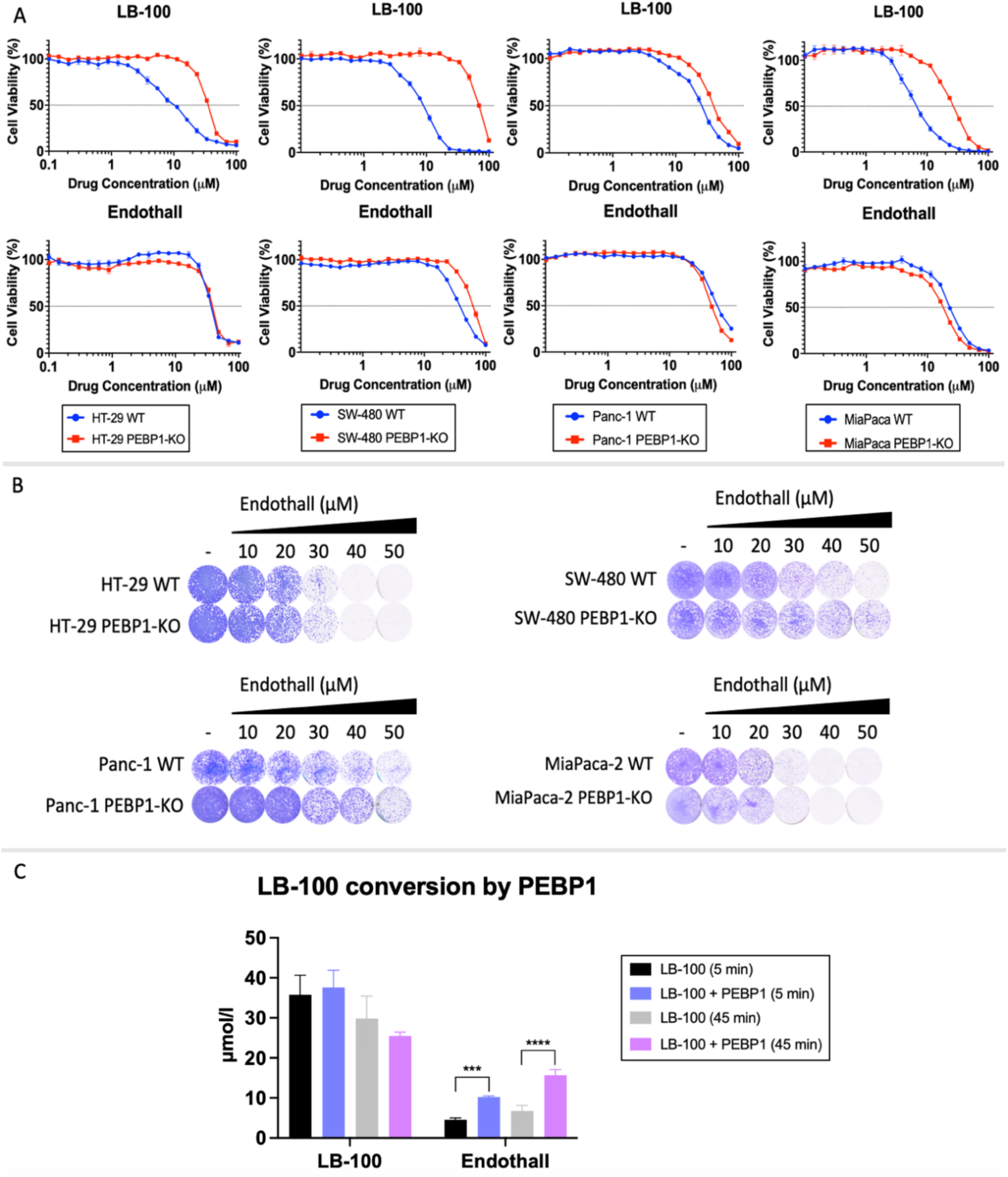
(A) Dose-response curves comparing the effect of the prodrug LB-100 and its active metabolite endothall in the presence or absence of PEBP1 in CRC and PC models. The values were normalized to PAO and DMSO, which were used as a positive control (0% cell viability) and negative control (100% cell viability), respectively. Mean values and their SD are shown in the graphs. (B) Long-term viability assay of the HT-29, SW-480, Panc-1, and MiaPaca-2 WT and PEBP1-KO cells treated with increasing range concentrations of endothall. The assay was terminated when the untreated group reached confluence. (C) Mass-spectrometry analysis of LB-100 conversion by PEBP1. LB-100 was used as a negative control. The quantity of the prodrug LB-100 and the active metabolite endothall were measured in both conditions. Two different time points of the reaction were included 5 and 45 minutes. The mean and the SD of the values are exhibited. Unpaired parametric one-way ANOVA was used to calculate the significance. *** for p≤0.001 and **** for p≤0.0001 significant level.

To further solidify this notion, we conducted long-term viability assays treating the CRC and PC cells with endothall. In this type of experiment, the drug was refreshed several times and the assays were terminated when the control groups reached confluence. Therefore, this method allowed for better observation of the drug’s effects compared to dose-response curves in which the cells were treated once, and the drug was not refreshed. The staining of the viability assay showed that the toxicity of endothall is not affected by the absence of PEBP1 (Figure 4B), which aligned with the results of the dose-response curves. Our data suggested that PEBP1 plays a critical role in the activation of the prodrug, as the cells were able to effectively respond to the active metabolite without the presence of the gene. Altogether, these findings support that deletion of PEBP1 confers resistance to LB-100 by reducing the conversion into endothall.

However, the above results only constitute indications of the hydrolase activity of PEBP1, as there are many other factors in the cells that have a hydrolase effect [45]. Therefore, we cannot conclude that the enzymatic reaction is performed by PEBP1. Several scenarios could explain the observation of low-PEBP1 mediated resistance to LB-100 and not to endothall. For instance, it could be possible that PEBP1 is part of a complex of proteins with catalytic activity or modulates the function of another enzymatic protein.

To test directly our hypothesis, we measured the cell-free conversion of LB-100 into endothall in the presence or absence of PEBP1 at two different time points (after 5 and 45 minutes). If PEBP1 would mediate the conversion, the final amount of endothall produced should be higher in the presence of the protein. The mass-spec analysis revealed that a small part of LB-100 was being converted to endothall in the PEBP1-free environment. However, the production of endothall was significantly increased in the presence of PEBP1 (Figure 4C). More specifically, the quantity of the prodrug was doubled by the contribution of the protein in comparison with the control group. These data were consistent in both time points. Altogether, these findings support the contribution of PEBP1 to the conversion of LB-100 into endothall and provide mechanistic insight into the resistance mediated by PEBP1 depletion.

## Discussion

Drug resistance is a significant challenge that hinders the effective treatment of cancer [6, 11]. Unfortunately, even patients who initially respond well to therapy may later develop resistance, undermining the effectiveness of the treatment and resulting in cancer progression and decreased overall survival [10, 11]. In this regard “paradoxical” hyperactivation of oncogenic signaling could be a potential therapeutic approach to over-expose cancer cells to stress and promote cell death when combined with stress response inhibitors [13]. Our recent research has identified LB-100 and adavosertib as promising molecules for combination therapy against multiple cancer models [23]. Despite the encouraging results obtained during this work, resistance will likely be a challenge for this proposed therapy, as observed for other therapies. The mechanisms of resistance to cancer therapies are complex and heterogeneous and their elucidation is essential for identifying potential drug targets and developing effective strategies [46]. In the present study, we examined how PEBP1 depletion can drive resistance to LB-100.

On the basis of PEBP1’s function in inhibiting signal transduction, it is postulated that this protein has a tumor-suppressive role [36]. Therefore, the decrease in PEBP1 expression was an unexpected mechanism underlying resistance to LB-100 [23]. The localization of PEBP1 in close proximity to the cellular membrane [47] and its influence on LB-100 toxicity and function suggested that PEBP1 can be involved in drug intake. Nonetheless, our experimental investigation using both water-soluble (LB-100) and lipid-soluble (LB-102) derivatives of endothall [42] demonstrated that this is not the case. Moreover, further analysis showed that depletion of PEBP1 decreased the toxicity of the prodrug LB-100 but not that of the active metabolite endothall, indicating that it may be involved in the processing of the drug intracellularly. These results in combination with the findings of Kazui et al (2016) proposed that PEBP1 might possess potent enzymatic activity and participate in drug metabolism [38], which could account for the resistance acquired by low-PEBP1 levels. The cell-free measurement of LB-100 and endothall in the presence of PEBP1 effectively validated our hypothesis that PEBP1 mediates the LB-100 conversion. It is possible that the hydrolase function of PEBP1 extends also in the metabolism of other drugs and could explain how its reduced expression has been linked to unfavorable prognosis and poor survival rates in cancer patients [33, 37]. These observations could relate the resistance developed by PEBP1 absence to the inefficiency of treatment.

In pursuit of this objective, it is feasible to identify the specific enzymatic reaction responsible for the release of endothall and explore the possibility of other drugs being activated through the same mechanism. Furthermore, we could examine the differences in hydrolase expression between the models being used and establish a correlation with the response to treatments. Additionally, genetic screens for candidates that are synthetic lethal with PEBP1-depletion or can revert the resistance mediated by the downregulation of PEBP1, will propose drug combinations that can mitigate or even benefit from the absence of the protein.

PEBP1 could become a potential biomarker for guiding the selection of specific targeted therapies and enhancing their translational efficacy. In targeted and other therapies, it is crucial to develop predictive biomarkers of sensitivity or resistance enabling clinicians to tailor treatment plans accordingly, by identifying patients likely to benefit from a particular medication and avoiding exposure to treatments that will not be effective [48]. Overall, identifying biomarkers of resistance/sensitivity is critical to improving patient outcomes and understanding tumor biology [10, 13, 48].

## Acknowledgments

We thank Emma van Dijk and Lieke Wilms for their help in performing some experiments and Patrick Celie for making recombinant PEBP1 protein. This work was supported in part by a grant from Lixte Biotechnology.

